# Disassembly of actin and keratin networks by Aurora B kinase at the midplane of cleaving *Xenopus laevis* eggs

**DOI:** 10.1101/513200

**Authors:** Christine M. Field, James F. Pelletier, Timothy J. Mitchison

## Abstract

We investigated how bulk cytoplasm prepares for cytokinesis in *Xenopus laevis* eggs, which are large, rapidly dividing cells. The egg midplane is demarcated by Chromosomal Passenger Complex (CPC) localized on microtubule bundles between asters. Using an extract system and intact eggs we found that local kinase activity of the AURKB subunit of the CPC caused disassembly of F-actin and keratin between asters, and local softening of the cytoplasm as assayed by flow patterns. Beads coated with active CPC mimicked aster boundaries and caused AURKB-dependent disassembly of F-actin and keratin that propagated ~40 μm without microtubules, and much farther with microtubules present, due to CPC auto-activation. We propose that active CPC at aster boundaries locally reduces cytoplasmic stiffness by disassembling actin and keratin networks. This may help sister centrosomes move apart after mitosis, prepare a soft path for furrow ingression and/or release G-actin to build the furrow cortex.

## Introduction

Cytokinesis in animal cells requires large scale re-organization of the cytoplasm involving essentially every cytoskeletal and membrane system. Most studies of cytokinesis mechanics focus on the cortex, where local actomyosin contraction promotes cleavage furrow ingression. The internal cytoplasm also has to re-organize to allow furrow ingression, but it is unclear to what extent this occurs by passive deformation in response to forces generated by the ingressing furrow, versus active reorganization by furrow-independent mechanisms.

Eggs of the frog *Xenopus laevis* are well suited for investigating the organization of internal cytoplasm prior to cleavage. They are ~1.2mm in diameter, and the 1^st^ furrow has to ingress hundreds of microns over tens of minutes. Although living eggs are opaque, an actin-intact egg extract system allows cell-free imaging of the cytoskeletal assemblies that control cytokinesis (Nguyen and Groen et al., 2014). The 1^st^ furrow cuts through the midplane of the egg, starting at the animal pole. This path is specified by an egg-spanning plane of microtubule bundles coated with cytokinesis signaling protein complexes CPC and Centralspindlin (Field et al., 2015). This plane initiates during anaphase at the position previous occupied by the metaphase plate, stimulated by proximity to chromatin. It then expands outwards at the boundary between the sister microtubule asters that grow from the poles of the mitotic spindle. Auto-activation of the CPC on stable microtubule bundles creates a positive feedback loop that propagates the CPC signal outwards from anaphase chromatin to the plasma membrane (Mitchison and Field, 2017). Once at the cortex, CPC-coated microtubule bundles trigger furrow assembly and ingression, which involves stimulation of actomyosin assembly (Basant and Glotzer, 2018). Here, we investigate their influence on actin and keratin networks in bulk cytoplasm far from the cortex.

The internal cytoplasm of *Xenopus* eggs contains F-actin and keratin networks with the potential to impede cleavage furrow ingression. From proteomic data, estimated concentrations in eggs are 14 μM actin, 700 nM keratin-8 and little vimentin or other intermediate filaments (Wuhr et al., 2014). Based on extract experiments, approximately 50% of the actin polymerizes into a dense network of bundles throughout the egg during both mitosis and interphase (Rosenblatt et al., 1995; Field et al., 2011). Keratin assembly increases after fertilization, presumably in response to decreased Cdk1 activity, and cleaving blastomeres contain an organized keratin filament array that is enriched on the cortex (Klymkowsky et al., 1987; Klymkowsky, 1995). The precise organization of keratin in the deep cytoplasm between microtubule asters has not been described.

The organization of actin and intermediate filaments during cytokinesis has been extensively investigated in smaller somatic cells. Actin polymerization locally increases at the furrow cortex during cytokinesis in response RhoA signaling (Basant and Glotzer, 2018). There has been less research on actin dynamics deeper in the cytoplasm, in part because cortical actomyosin is thought to dominate the mechanical behavior of small cells. Keratin and vimentin filaments are distributed throughout the cytoplasm in somatic cells. They partially depolymerize during mitosis, in response to phosphorylation by CDK1 (Izawa and Inagaki, 2006). AURKB, the kinase subunit of the CPC, was also implicated in depolymerizing vimentin during cytokinesis, and specific sites involved in this regulation were mapped (Goto et al., 2003). Keratin is also controlled by CDK1 and AURKB phosphorylation during cell division (Inaba et al., 2018). The keratin network appears to become globally disassembled at the onset of mitosis, then starts reforming during cytokinesis (Lane et al., 1982). One plausible model is that CDK1 phosphorylation disrupts the keratin network, and AURKB phosphorylation prevents its premature re-assembly in the furrow.

Here, we mainly use an egg extract system to investigate how boundaries between asters influence cytoplasmic organization. We find that CPC localized at aster boundaries locally disassembles actin and keratin networks by mechanisms that depend on AURKB kinase activity, and this leads to local softening of the cytoplasm between asters. We then recapitulate actin melting activity using CPC artificially recruited and activated on beads, and show that keratin disassembles ahead of the furrow in dividing eggs. We propose the F-actin and keratin disassembly activities of the CPC prepare egg cytoplasm for furrow ingression by clearing a soft path in advance of the furrow.

## Results

### AURKB-dependent disassembly of F-actin bundles at boundaries between asters

Figure 1A and Movies 1A, B and C show a typical experiment in actin-intact egg extract where interphase was triggered by calcium addition. Microtubule asters were nucleated from beads coated with anti-AURKA IgG (Tsai and Zheng, 2005), F-actin was visualized with Lifeact-GFP and CPC with labeled anti-INCENP IgG. As soon as we initiated imaging, typically 1-3min after warming the extract from 0° to 20°C, F-actin was present as a dense network of relatively homogenous bundles interspersed with occasional comet tails. In the vicinity of microtubule organizing centers the actin bundles became partially entrained to the radial array of microtubules at early time points (Fig 1A 8, 18min). After CPC was recruited to aster boundaries the network of actin bundles locally disappeared, starting at the line of CPC recruitment, and spreading outwards over tens of μm in tens of min. By 60min, F-actin bundles were mostly restricted to islands centered on MTOCs, separated by gaps ~100μm wide centered on CPC-coated anti-parallel microtubule bundles. This organization persisted for 10s of minutes. Loss of F-actin bundles at CPC-positive aster boundaries was highly reproducible. Every CPC-positive aster boundary exhibited F-actin depletion across >100 fields in >30 batches of extract.

F-actin comet tails, presumably nucleated by Arp2/3 on endosomes (Taunton et al., 2000), were not inhibited at aster boundaries. The arrowheads and 3x inserts in Fig1A highlight individual F-actin comet tails in boundary regions where the main actin network is disassembled. See Movie 1C. It was difficult to compare the density of comet tails in cleared vs non-cleared regions, but their shape brightness and movement rate were qualitatively similar. The presence of normal-looking comet tails at aster boundaries depleted of actin bundles suggests that spatial regulation does not involve a complete block to actin polymerization, and might be based on negative regulation of a formin-dependent nucleation mechanism.

**Figure 1.**
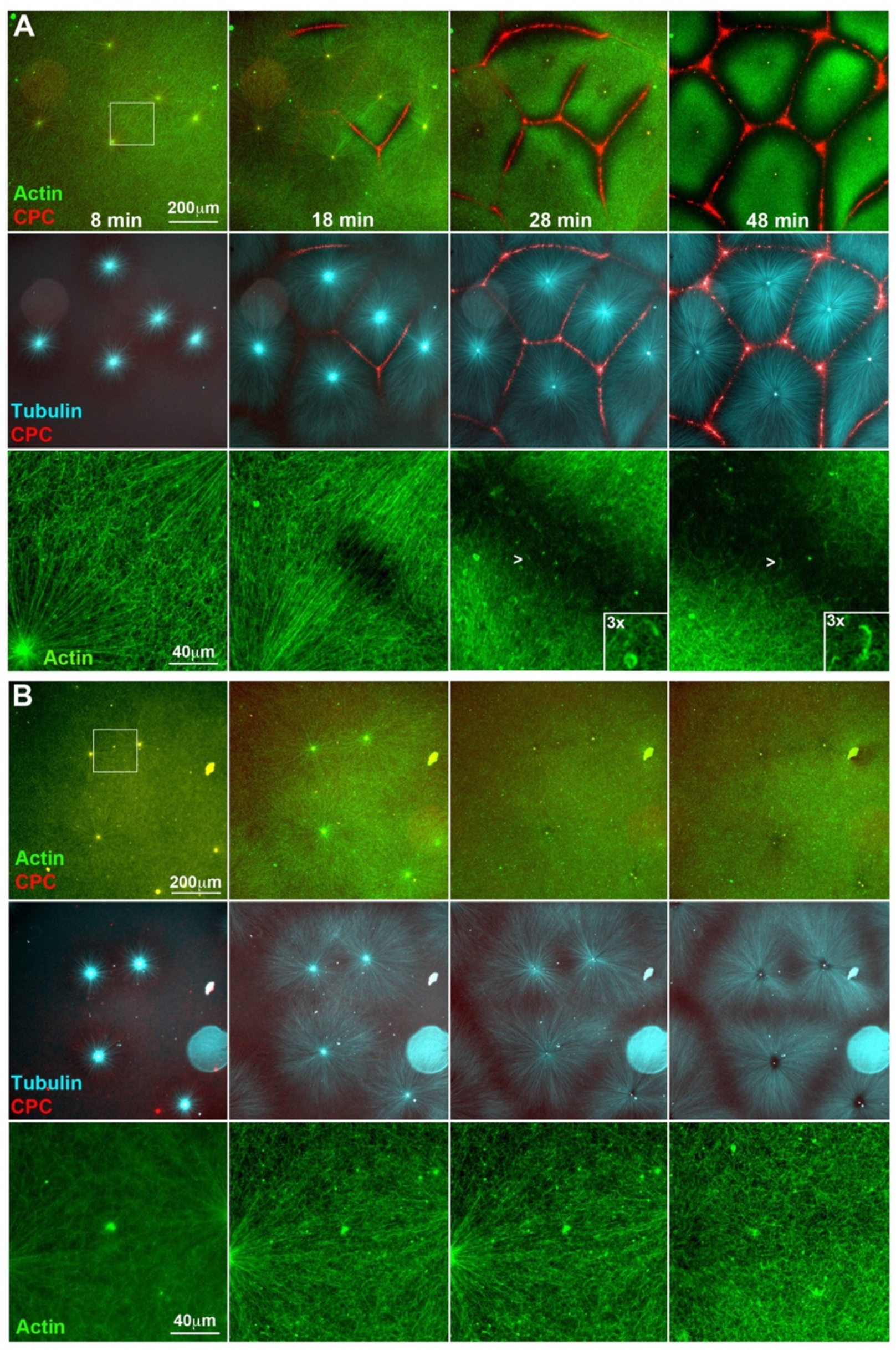
AURKB-dependent disassembly of F-actin at boundaries between asters in egg extract. **A)** Control reaction containing probes for F-actin (GFP-Lifeact), CPC (A647-anti-INCENP IgG) and microtubules (Tau peptide-mCherry) imaged by wide-field epifluorescence with a 20x objective. Note recruitment of CPC to anti-parallel microtubule bundles at boundaries between asters, and loss of F-actin bundles where CPC is recruited. The third row is higher magnification views of the region boxed in the top left panel. Note residual F-actin comet tails in the cleared region (chevrons, and 3x inserts). See Movies 1A, B and C. **B)** Reaction containing 40 μM barasertib, a selective AURKB inhibitor. Asters grow out and contact each other, but CPC recruitment is blocked, and F-actin bundles are not disassembled at aster boundaries. The third row is higher magnification views of the region boxed in the top left panel. See Movies 2A and B.

To test if disassembly of F-actin bundles at the boundary between asters depended on AURKB kinase activity we added the selective small molecule AURKB inhibitor barasertib (Mortlock et al. 2007). This reagent completely blocked CPC recruitment (Fig 1B) and actin clearing, and had less effect on microtubule organization within asters than other AURKB inhibitors tested (ZM447439, hesperadin). AURKB inhibition slowed initial nucleation of asters, but large aster still grew out after a lag of several minutes. AURKB is required for formation of sharp microtubule boundaries between asters with little interpenetration of growing +TIPs from neighboring asters (Nguyen and Groen et al., 2014). With barasertib, most asters did not form sharp microtubule boundaries, however some asters formed less robust microtubule boundaries at later times, likely because the “anti-parallel pruning” activity of PRC1E + Kif4A does not completely depend on AURKB (Nguyen et al., 2018). F-actin clearing at aster boundaries was completely inhibited by barasertib (Fig 1B, Movies 2A and B). F-actin appeared uniform across the whole field except for some initial entrainment to microtubules in the vicinity of organizing centers (Fig 1B).

### AURKB-dependent disassembly of keratin at boundaries between asters

Fig 2A shows keratin organization as asters meet, using labeled anti-keratin IgG for visualization as previously reported (Weber and Bement, 2002). Fig 2A and 2C are taken from an image sequence collected in parallel with Fig 1A and B, so conditions and timing can be directly compared. EB1-GFP was used to visualize microtubule growth in this experiment Network organization of keratin was disrupted at the boundary between asters, spatially coincident with CPC recruitment (Fig 2A). Keratin disruption was slower than F-actin, and extended less far from the aster boundary, perhaps reflecting the less dynamic nature of keratin filaments. As with F-actin depletion, disruption of keratin by the CPC eventually led to formation of keratin network islands centered on MTOCs (Fig 2A). We often noted accumulation of disassembly keratin fragments at the center of aster boundaries (Fig 2A 3x inset). We suspect this is due to plus end-directed transport of keratin fragments that accumulate when the network is partially dissolved by AURKB activity. Movies 3A and B show a different control experiment where keratin disassembly between asters is clearly visualized.

The precise morphology of keratin disassembly at aster boundaries depended on the relative kinetics of mitotic exit vs boundary formation. In mitotic extract, where CDK1 activity is high, the keratin signal exhibited little spatial contrast, suggesting keratin was organized into polymers too small to resolve (not shown). After triggering the extract to enter interphase, the keratin image gradually coarsened over ~30min, evolving first into homogenously-distributed particles, and then into a continuous filament network whose contrast increased over time (see panels at the bottom of Fig 2C). This gradual coarsening, which was similar with and without AURKB inhibitor, provided visual evidence for progressive assembly of keratin into a continuous network when CDK1 activity drops. When the aster density was high, CPC-positive boundaries formed before keratin had fully assembled into networks. In this case, keratin networks never formed in the boundary region, and keratin fragments tended to accumulate at the center of the boundary (Fig 2A). When aster boundaries formed late in the reaction, an extended keratin network was already present at the time of CPC recruitment. In this case the pre-formed keratin network locally disassembled when CPC was recruited to microtubule bundles at the aster boundary (Fig 2B, Movie 3C).

Keratin disassembly at aster boundaries depended on AURKB activity, since it was blocked by barasertib (Fig 2C, Movie 4). Keratin networks were still reduced around MTOCs, and in some cases were slightly enriched at aster boundaries. Both effects could be due to kinesin-mediated transport. To quantify keratin disassembly, we scored 28 asters on 4 coverslips under control conditions at 45 min. Out of 58 boundaries that were positive for CPC, 58 (100%) showed evidence of keratin disassembly. On 4 coverslips run in parallel with 40 μM barasertib, we scored 18 aster boundaries. Zero boundaries were positive for CPC and zero showed evidence of keratin disassembly. We did sometimes observe areas of low keratin density which corresponded to organelle aggregates.

**Figure 2.**
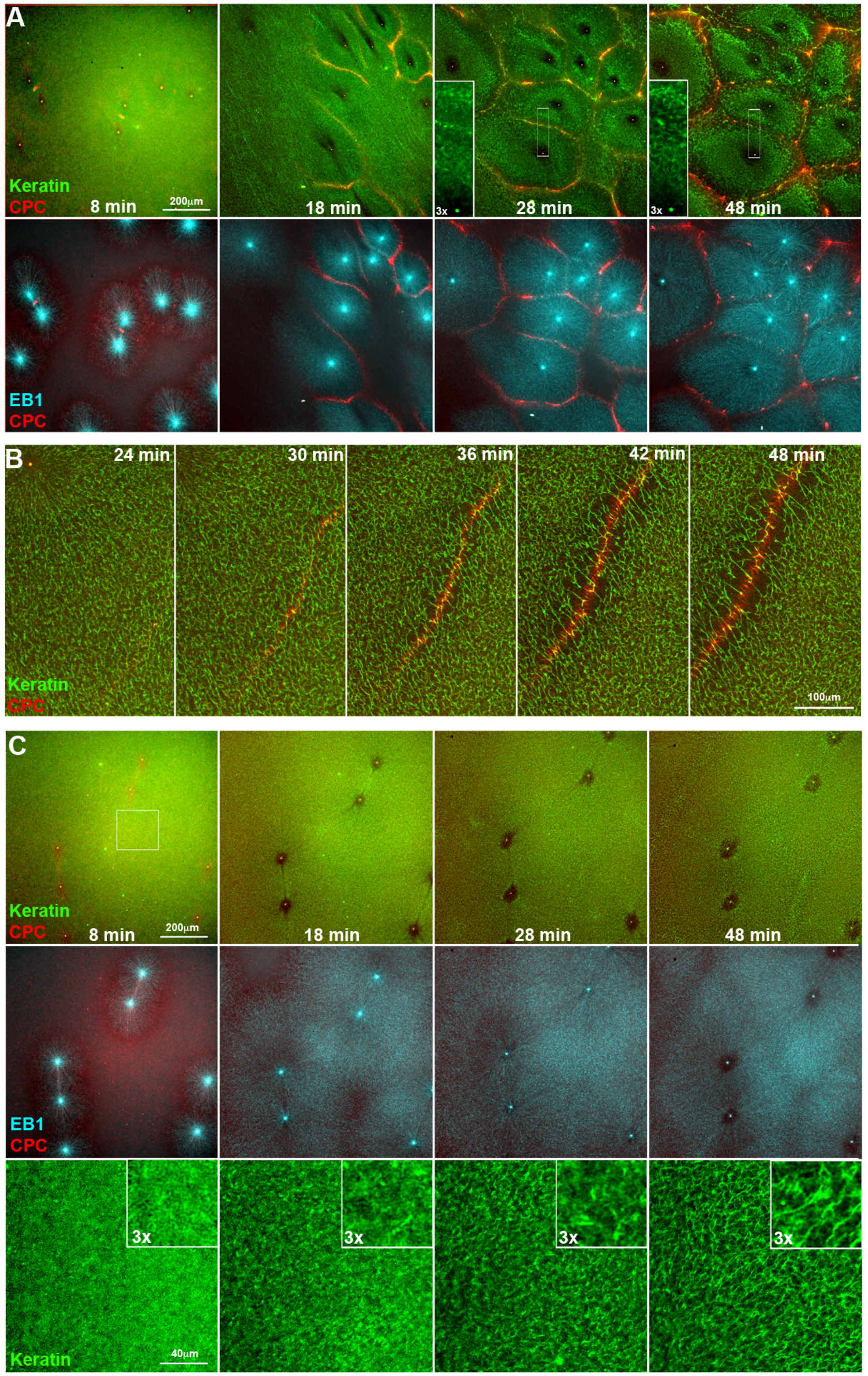
AURKB-dependent disassembly of keratin at boundaries between asters in egg extract. Panels A and C are from reactions run in parallel with Fig 1 using multiposition time lapse, so timing can be directly compared. **A)** Control reaction containing probes for Keratin (A568-anti-keratin IgG), CPC (A647-anti-INCENP IgG) and growing microtubule plus ends (EB1-GFP). Note partial disassembly of keratin at boundaries between asters where CPC is recruited. Keratin aggregates transiently accumulated at boundary lines, which is noticeable at 28 min in this example. Movies 3A and B show similar data from a different experiment. **B)** Images from an experiment similar to A) but run on a different day. This example was chosen to highlight a boundary zone that formed late in the reaction, after the keratin network was already established. Note disruption of the network along the line of CPC recruitment. Bulk flow caused the whole network to move leftwards and upwards in this time series. See Movie 3C. **C)** Parallel reaction to A) containing 40μM barasertib. See Movie 4. CPC recruitment and keratin disassembly at asters boundaries are both blocked. Keratin clearing in a circular zone near MTOCs, presumably caused by transport and/or outwards microtubule sliding, is similar to the control reaction in A). The bottom row shows higher magnification views of the boxed region in the top left panel. Note gradual evolution of the keratin signal from homogeneous aggregates into a network of connected bundles. Similar time-dependent assembly occurred in control reactions, and is presumably due to progressive assembly following loss of mitotic Cdk1 activity.

The labeled anti-INCENP IgG probe used to visualize the CPC in Figures 1&2 promotes auto-activation of the CPC (Field et al., 2015). This was convenient for observation of F-actin and keratin disruption because every aster boundary recruited CPC within a few minutes of the asters interacting. To test if actin and keratin disruption depended on use of an activating CPC probe we substituted a GFP fusion of the Dasra subunit of CPC, which exchanges into the complex, and does not stimulate CPC auto-activation (Nguyen et al., 2014). Using this probe, some aster boundaries recruit CPC and others in the same field do not, dependent on the initial distance between MTOCs (Field et al., 2015). We imaged aster assembly reactions using Dasra-GFP and analyzed data at a timepoint where ~50% of aster boundaries had recruited CPC. For quantification, we used a Voronoi diagram to define aster boundaries in the microtubule channel, then scored CPC recruitment and actin clearing at these boundaries in the other channels. 92 aster boundaries were scored on 3 coverslips. Of these, 43 were CPC positive and disassembled actin. 45 were CPC negative and did not clear actin. 1 was CPC positive and did not clear actin, and 3 appeared to clear actin but were not CPC positive. Thus, 88/92 (96%) matched our prediction that CPC localization correlates with actin clearing. These data confirmed that actin clearing occurs close to CPC-coated microtubule bundles with a non-activating CPC probe. By visual inspection, the kinetics and extent of actin clearing were independent of the CPC probe used.

### Boundary regions are less gelled than aster centers

Disruption of F-actin and keratin networks at aster boundaries is likely to cause a change in local mechanics of the cytoplasm. We probed this by measuring cytoplasmic flows in response to pressure gradients. Flow was induced by adding and removing a small weight from the top coverslip and imaged in the DIC channel as coherent movement of small particles that correspond to mitochondria and other vesicles. When flow was induced after assembly of CPC-positive aster boundaries it tended to occur preferentially at boundary regions, while particles closer to aster centers remained stationary, or moved slowly on radial tracks. Fig 3 and Movie 5 show a typical example. Particle Image Velocimetry (PIV) was used to track rapid particle movement in the DIC channel. Note preferential flow in channels demarcated by loss of F-actin (Fig 3C). To check if this was reproducible, we visually scored 21 fields on two separate days (two different extracts) for evidence of flow following transient compression of the coverslip. We observed flow preferentially through aster boundaries in 18 examples, no clear flow in 3, and zero examples with flow through aster centers. In parallel reactions containing barasertib to prevent CPC recruitment and F-actin clearing we observed zero examples of locally coherent flow between asters in 15 examples. In most cases compressing the coverslip caused the whole field to displace without preferential flow channels, in a few cases we observed rips and flows in the gel in random locations. We conclude that the boundary between asters provides a soft path for flow of cytoplasm and formation of this path depends on local melting of cytoskeleton networks by AURKB activity.

**Figure 3.**
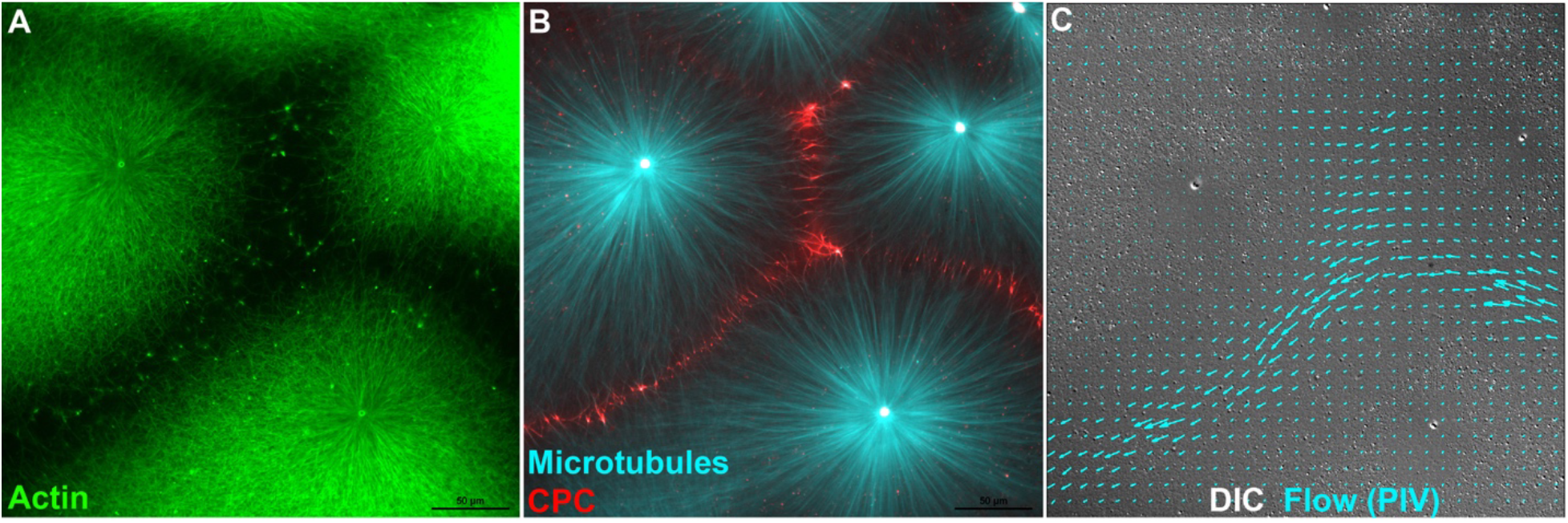
Low resistance to pressure induced flow between asters. **A,B)** Fluorescence images from an aster assembly reaction similar to Fig 1. t=25min. **C)** DIC image of the same field, taken from a movie captured after the coverslip was transiently compressed to generate flow (See Movie 5). Particle Image Velocimetry (PIV) analysis of two sequential frames from the end of the movie was used to measure rapid particle movement indicative of flow. Blue arrows indicate movement direction and velocity from PIV. Flow occurred preferentially through the F-actin free channels between asters. Peak flow rate was ~7μm/sec in this example. Actin (Lifeact-GFP), Tubulin (Tau peptide mCherry) and CPC (anti-INCENP-Alexa-647.

### Disassembly of F-actin around CPC-coated beads

The CPC-coated microtubules bundles that assemble between asters could interact with F-actin in multiple ways, e.g. via motor proteins or MAPs. To test if local concentrations of active CPC in the absence of microtubules were sufficient to disrupt F-actin and keratin networks we prepared beads coated with anti-INCENP IgG, which is known to cause auto-activation of bound CPC (Sampath et al., 2004; Field et al., 2015). To determine what proteins bound to these beads we performed mass spectrometry after incubating in extract under various conditions (full data available on request). In 3 experiments all 4 CPC subunits were among the most abundant peptides bound to anti-INCENP beads. To a more variable extent we observed known CPC interaction partners Kif4A, the egg homolog of Kif20A (aka MKLP2) and NPM1/2. Actin, keratin and the actin-binding protein filamin were bound to variable extents, but these are standard contaminants in our IP-MS experiments from egg extract, and thus hard to interpret. We did not observe recruitment of known actin or microtubule depolymerizing factors.

To test if beads coated with active CPC could locally disassembly F-actin networks we started by imaging in the presence of nocodazole (Fig 4 A, B). This simplified the reaction by preventing microtubule-based auto-amplification of CPC activity (Sampath et al., 2004). We used 7μm beads to maximize local effects of active CPC. We observed reproducible clearing of F-actin extending ~20μm away from each bead by eye, and remaining roughly constant in diameter over time (Fig 4A). Clearing was not observed around control IgG beads (not shown) or when AURKB activity was inhibited with barasertib (Fig 4B). To quantify F-actin clearing, and its AURKB-dependence, we first scored beads by eye. In one experiment 30/31 anti-INCENP beads vs 0/59 beads coated with control IgG exhibited local clearing of F-actin at 45min. In a different experiment 22/23 anti-INCENP beads vs 0/23 incubated in parallel with barasertib exhibited F-actin clearing at 45min. We next quantified the spatial extent of F-actin clearing by computing radial F-actin intensity profiles with respect to distance from the beads (Fig 4C). Clearing was detectable extending up to 40 μm from anti-INCENP beads with a half-maximal distance of ~15μm, and was blocked by barasertib. We interpret these data as revealing a reaction-diffusion system where an unknown actin regulator visits the beads, becomes phosphorylated by AURKB, diffuses away where it acts to disassemble F-actin. At the same time, it is dephosphorylated and inactivated by soluble phosphatases resulting in a gradient of activity. Keratin dissolution was weaker in assays containing nocodazole, and was observed around 7μm anti-INCENP beads in only some experiments.

Cytoskeleton clearing behavior around anti-INCENP beads was much more extensive when nocodazole was omitted (Fig 4 D, G, H and Movie 6). In this case standard 2.8μm magnetic beads caused reliable F-actin disassembly, and these were used for all reactions. With microtubules present we observed dramatic disassembly of F-actin around 2.8μm anti-INCENP beads (Fig 4 D, H), and also partial disruption of keratin (Fig 4D), partial loss of microtubules (Fig 4 D, H), and complete loss of EB1 comets (Fig 4G). F-actin was disassembled more completely, and over much larger distances, compared to reactions without microtubules (compare panels D&F). The radius of zones with disassembled F-actin and keratin increased with time from ~50μm to 200μm or more. Inhibition of all cytoskeletal networks around beads was AURKB activity-dependent since it was not observed around beads coated with control IgG (not shown), or when an AURKB inhibitor was added (Fig 4E). To quantify AURKB dependence we scored F-actin, tubulin and keratin clearing by eye in one typical experiment. Out of 17 beads scored in the control reaction, the numbers that exhibited clearing for each filament system at 60 min were: F-actin 17, microtubules 16, keratin 14. In a parallel reaction containing barasertib 31 beads were scored, and F-actin disassembled around 0/31 beads, microtubules around 1/31 beads, and keratin around 0/31 beads.

**Figure 4.**
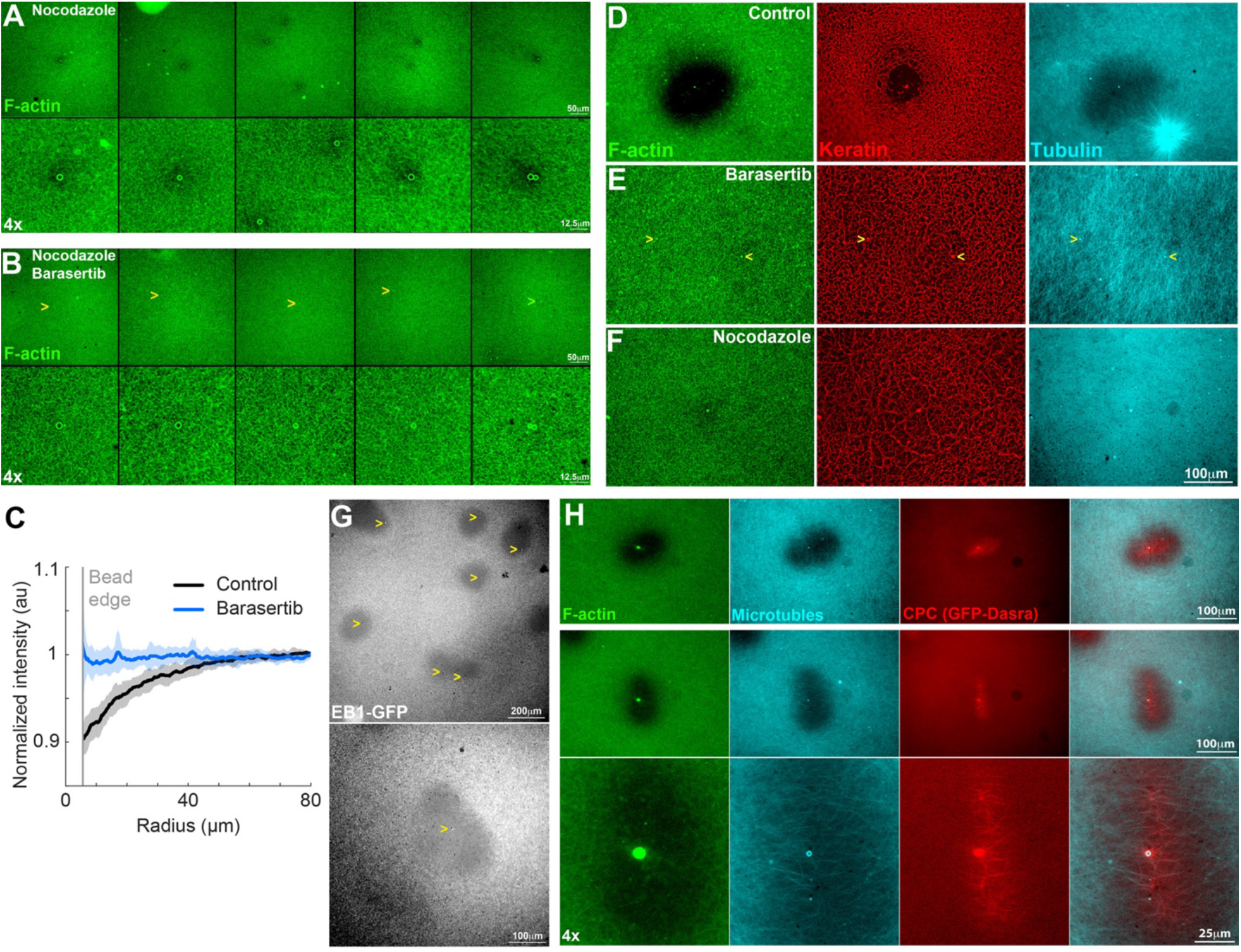
Inhibition of cytoskeleton assembly around CPC-coated beads. 7μm beads (panels A,B) or 2.8μm beads (panels C-H) coated with anti-INCENP IgG were used to recruit and activate endogenous CPC in extract. 40μM nocodazole and/or 40μM barsertib were included in reactions as indicated to inhibit microtubules and/or AURKB. Yellow chevrons indicate CPC beads in panels where they are difficult to see. Actin (Lifeact-GFP), keratin (A568-anti-keratin IgG), Tubulin (directly labeled with ALEXA-647 NHS ester), CPC (DasraA GFP) and growing microtubule plus ends (EB1-GFP). **A)** Partial F-actin clearing around 7μm CPC beads in the absence of microtubules visualized with Lifeact-GFP. t=45min. Partially disassembled zones were ~40μm in diameter. **B)** AURKB activity is required for partial F-actin clearing around 7μm CPC beads. t=45min. Panels A, B are taken from parallel reactions. Note no loss of F-actin around beads. **C)** Quantification of F-actin clearing around 7μm CPC beads in the absence of microtubules using radial intensity plots. 15 beads per condition. Dark lines show average radial intensity values, pale regions show the range. Linescans were truncated below 7μm to avoid the strong bead edge signal. **D)** Complete disassembly of F-actin, partial disassembly of keratin, and partial inhibition of microtubules around 2.8μm CPC beads when microtubules are present. t=60min. Disassembled zones were initially ~50μm in diameter, and grew progressively to 200μm or more. See Movie 6. **E)** AURKB activity was required for the clearing shown in D. t=60min. **F)** Microtubules were required for the keratin disassembly and microtubule clearing shown in D. t=60min. Partial F-actin clearing, similar to panel A, is still visible in this image, but this was variable with 2.8μm CPC beads. Panels D, E, F are taken from parallel reactions. **G)** Complete EB1 comet clearing around CPC beads. Conditions similar to D, t= 60min. **H)** CPC recruitment around CPC-coated beads. Conditions similar to D, t= 60min. Note that CPC beads are surrounded by a network of CPC-coated microtubule bundles which assemble by capture of spontaneously nucleated microtubules. These amplify CPC recruitment and explain why F-actin is disassembled over much larger distances when microtubules are present.

Large, spreading zones of F-actin disassembly only initiated around anti-INCENP beads after incubation of squashes for >40min. Dynamic microtubules assemble spontaneously in interphase extract 20-40 min after triggering interphase by calcium addition, and rapidly increase in density (Ishihara et al., 2014). To understand how large zones of F-actin disassembly form we co-imaged microtubules, F-actin and Dasra-GFP, a non-activating CPC probe. Although the overall microtubule density was low in the inhibited zones, we observed formation of CPC-coated microtubule bundles in the vicinity of the beads at 60min. These bundles appeared disorganized in some cases, and aligned in parallel arrays in others (Fig 4H). Since the area around these bundles was completely devoid of EB1 comets (Fig 4G), these bundles must lack growing plus ends. We hypothesize these networks of non-dynamic, CPC-coated microtubule bundles form and grow by accretion of spontaneously nucleated microtubules from bulk solution. Once formed, they locally disrupt F-actin and keratin networks, and also inhibit growing microtubule ends. In these bead-triggered reactions, CPC-coated bundles form by aggregation of spontaneous microtubules accompanied by CPC binding and autophosphorylation (see discussion). This assembly mechanism differs from that of normal CPC-coated microtubule bundles, which form at aster boundaries when parallel microtubules from each aster grow into each other (Mitchison and Field 2017). However, we believe the CPC recruitment and auto-amplification mechanism is the same in both cases.

### Disruption of keratin organization ahead of the cleavage furrow in *Xenopus* eggs

To look for disruption of the cytoskeleton cleavage ahead of furrow ingression in eggs we fixed zygotes before and during 1^st^ cleavage (70-100min post fertilization) and imaged keratin organization by immunofluorescence (Fig 5). Despite considerable effort we unable to image F-actin filaments in fixed zygotes. We observed clear evidence for disassembly of keratin at the boundary between sister asters in zygotes fixed prior to cleavage initiation (Fig 5A) and ahead of ingressing furrows in zygotes fixed during cleavage (Fig 5B, C). Keratin disassembly was spatially coincident with CPC at the boundary between asters, as expected from extract observations. The presence of yolk platelets and lipid droplets made it difficult to visualize extended filaments, but the appearance of keratin at boundaries between asters was quite similar in fixed zygotes and the extract system. Disassembly occurred over a zone ~30μm wide centered on the aster boundary. In zygotes that had not yet initiated cleavage we usually observed a thin plane of keratin aggregates at the center of the boundary region (Fig 5A). These resembled accumulation of keratin aggregates at the center of the boundary region in extract (Fig 2A). After furrow initiation (Fig 5B, C) the network organization of keratin was more pronounced throughout the zygote, and its disruption near CPC-coated microtubule bundles was more obvious. We conclude that keratin network disassembly occurs at the boundary between sister asters in zygotes, spatially coincident with CPC enrichment. Disassembly occurs hundreds of μm, and hundreds of secs, ahead of the advancing furrow.

**Figure 5.**
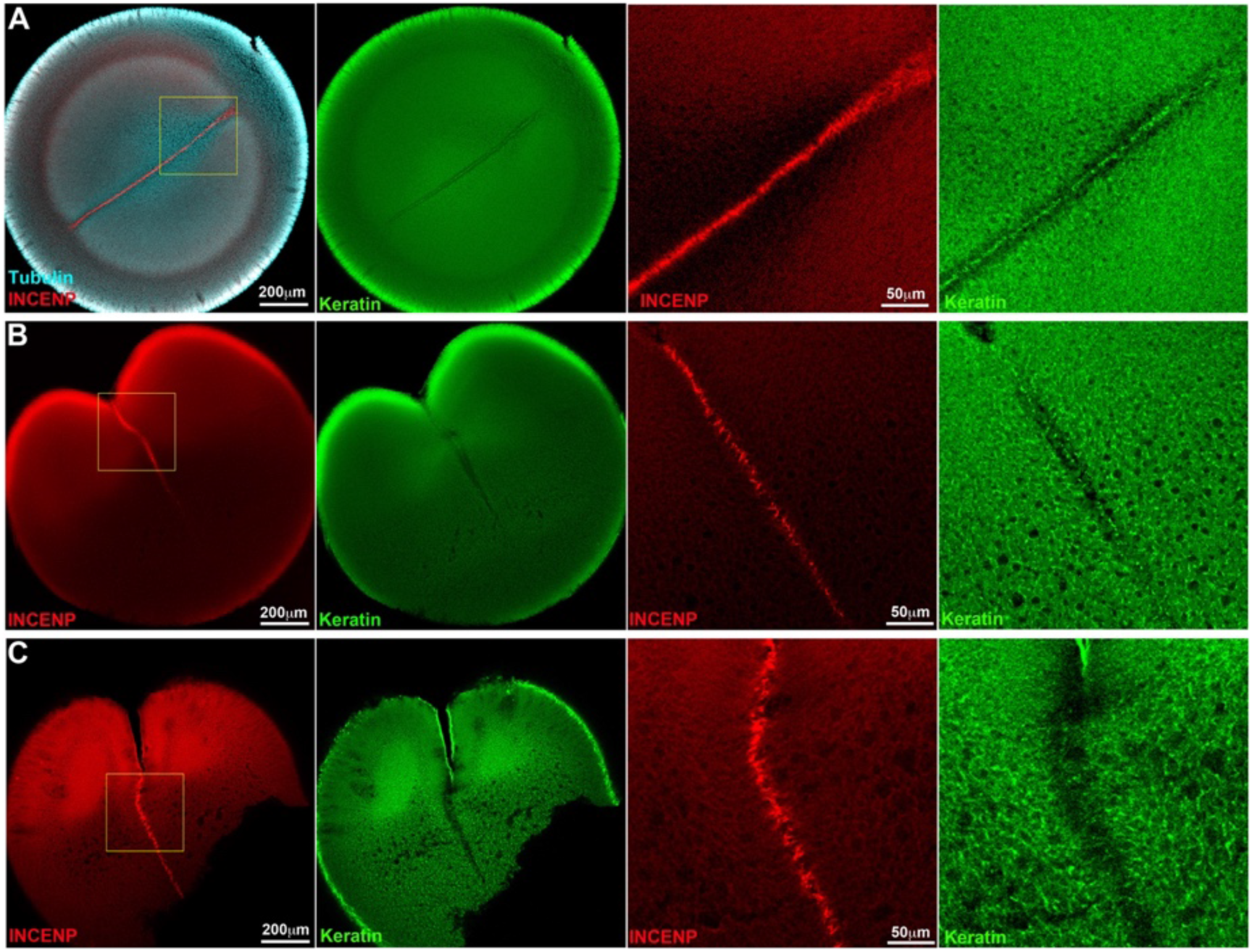
Keratin disassembly ahead of cleavage furrow ingression in fixed zygotes. Xenopus zygotes were fixed 70-100min post fertilization, bleached, stained and imaged by confocal microscopy in a clearing solvent. The second image pair in each row is a 4x view of the boxed region. Microtubules are poorly preserved by our fixation method once the furrow starts to ingress, and are only shown in A. **A)** 70min post fertilization, egg viewed from the animal pole. The asters have grown to ~65% of the egg radius, cleavage has not yet initiated. Keratin bundles are not yet pronounced in the bulk cytoplasm. Keratin appears partly disassembled in a ~30μm zone at the aster boundary. A line of keratin aggregates is present at the center of the boundary. These aster boundaries with a line of keratin fragments in the center resemble those seen in extract (Fig 2A) **B)** 90min post fertilization, animal pole is at top left. This zygote recently initiated cleavage. Keratin bundles are evident in the higher magnification view (which is a different focal plane). Note partial clearing, and accumulation of aggregates at the center of the aster boundary region, similar to A). **C)** 100min post fertilization, animal pole is at the top. The furrow has ingressed further than B), and keratin bundles are more pronounced away from the aster boundary. Large keratin bundles also accumulate at the cortex at this stage (not shown). Disassembly of the keratin network ahead of the ingressing furrow is evident, especially in the higher magnification view.

## Discussion

Figure 6A illustrates our current understanding of molecular pathways at the boundary between two asters. It is an extension of our previous model for autocatalytic spreading of CPC-coated microtubules at aster boundaries (red arrows). We previously hypothesized that autocatalytic spreading serves to relays information on the position of anaphase chromatin from the metaphase plate to the cortex to position furrows (Mitchison and Field, 2017). Our new observations strengthen the autocatalytic spreading model by showing that beads coated with active CPC activity induce CPC coating and bundling of randomly generated local microtubules, leading to spreading of CPC-positive islands in what appears to be an autocatalytic reaction (Fig 4D, H). Using F-actin melting in the absence of microtubules as an endogenous reporter of AURKB activity (Fig 4A), we could estimate the distance of action of a hypothetical reaction-diffusion system emanating from CPC beads. The half-distance of this gradient was ~15μm (Fig 4C). This value is important for building models of CPC spreading at aster boundaries. The CPC signal spreads between microtubule bundles normal to their axis, and a reaction-diffusion mechanism is one way this could occur. Another possible spreading mechanism would involve lateral fluctuation of microtubules, and spreading of the CPC-positive state by direct contact.

**Figure 6.**
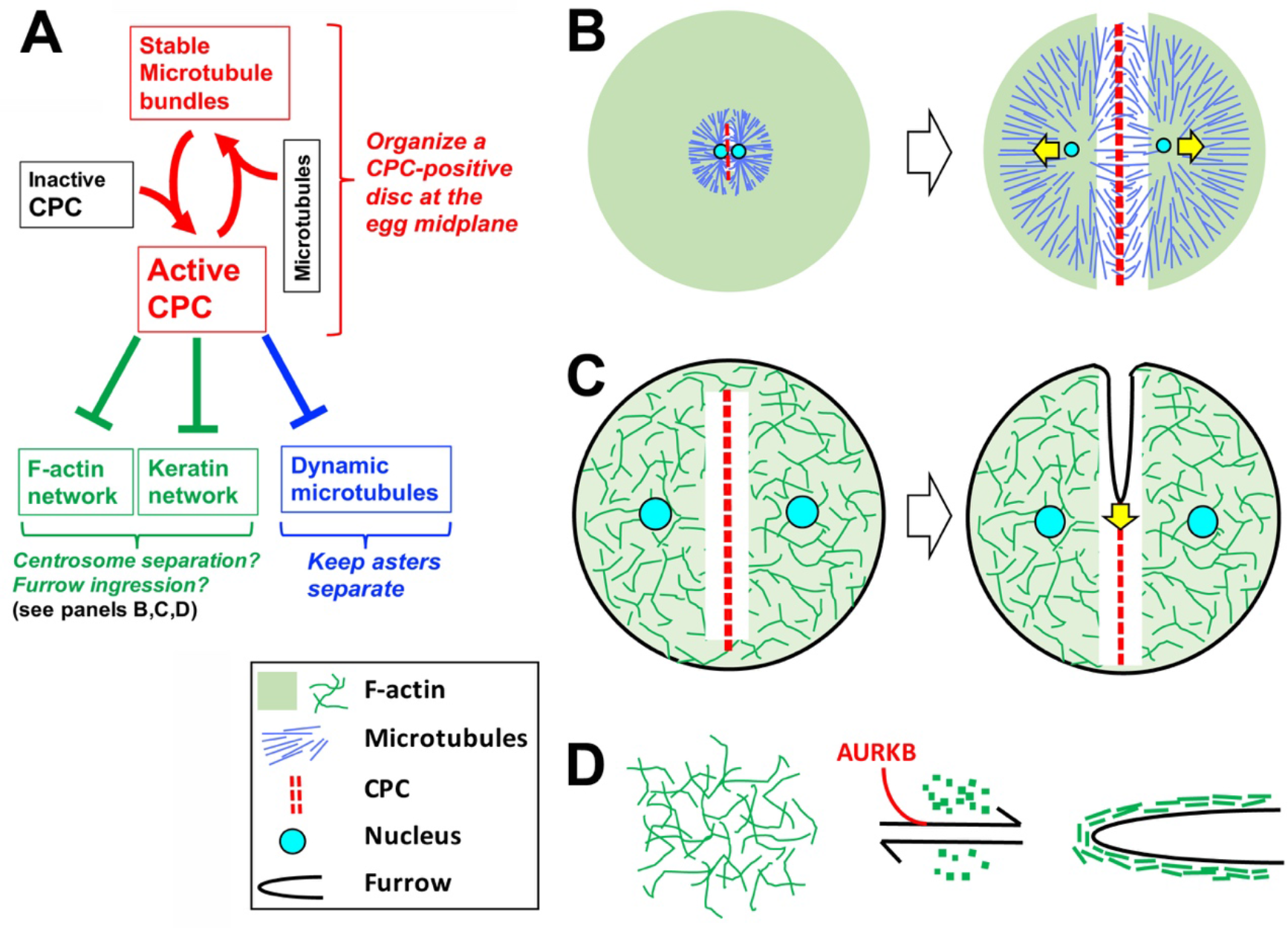
Model for CPC recruitment and functions at aster boundaries A) Summarizes molecular activities of the CPC, B-D) Illustrate possible functions of F-actin and keratin disassembly activities. **A)** Red arrows indicate a positive feedback loop that promotes recruitment and spreading of the CPC on microtubule bundles at aster boundaries (Sampath et al., 2004; Mitchison and Field, 2017). Green and blue negative arrows indicate AURKB-dependent inhibition of all three cytoskeletal systems, presumably via reaction-diffusion mechanisms. **B)** Disassembly of cytoskeletal networks between microtubule asters may help centrosomes and nuclei move apart following anaphase. **C)** Disassembly of cytoskeletal networks may help the furrow ingress. **D)** Disassembly of bulk F-actin may supply the ingressing plasma membrane with subunits to build the new cortex, in a model where different F-actin assemblies compete for subunits.

The mechanism by which AURKB locally disassembles F-actin (green inhibitory arrow in Fig 6A) is not known. AURKB stimulates furrowing when it reaches the cortex, and did not inhibit Arp2/3 comet tails at aster boundaries (Fig 1). Thus, it must negatively regulate bulk F-actin by a mechanism that is specific for some assemblies over others. One possibility is negative regulation of a formin that nucleates bulk cytoplasmic bundles, perhaps a homolog of FMN2 which nucleates bulk cytoplasmic actin in mouse eggs (Schuh and Ellenberg, 2008).

How AURKB destabilizes keratin filaments (green inhibitory arrow in Fig 6A) is also unknown, but in this case direct phosphorylation of keratin itself by AURKB is a likely model. Phosphorylation of intermediate filament polypeptides is a well-known mechanism for negatively regulating their assembly (Izawa and Inagaki, 2006), and AURKB sites on keratins 5&14 during cytokinesis were recently mapped (Inaba et al., 2018). Phosphorylation at related sites on keratin 8/18 is likely to directly inhibit keratin network assembly in the vicinity of CPC during cytokinesis in frog eggs.

AURKB regulation of microtubules at aster boundaries is complex, and involves both positive (red arrow in Fig 6A) and negative signals (blue inhibitory arrow in Fig 6A). This complexity reflects the central role of the CPC in generating stable microtubule bundles at the aster boundary, while at the same time destabilizing growing plus ends to keep the asters separated. We previously showed that asters are kept separate in part by the anti-parallel pruning activity of PRC1E and Kif4A (Nguyen et al., 2018). In our new experiments with CPC-coated beads, AURKB activity locally inhibited plus end growth of spatially disorganized microtubules formed by spontaneous nucleation, which is particularly evident in EB1 images (Fig 4G). This suggests anti-parallel organization is not required for inhibition of growing plus ends by AURKB activity. We suspect the CPC sends additional negative signals to keep asters separate during cytokinesis.

The biological functions of F-actin and keratin disassembly by the CPC are not known, though we did show that these activities cause a local softening or melting of the cytoplasm, such that hydrodynamic flow occurs much faster between asters than through them (Fig 3). We hypothesize three possible functions of this softening: helping sister centrosomes move apart after mitosis by softening the cytoplasm between them (Fig 6B), providing a soft path to help the furrow ingress (Fig 6C) and supplying G-actin subunits to build the cortex of the ingressing furrow and new plasma membrane (Fig 6D). The latter could be important given that different F-actin networks are known to compete for G-actin subunits (Zimmermann et al., 2017). Further experiments are required to evaluate these models, which are not mutually exclusive.

Finally, the egg extract system may hold lessons for organization of syncytial organisms. The frog zygote between anaphase and furrow ingression can be viewed as a temporary syncytium, where both nuclei are replicating. The aster-dependent organization of cytoplasm into distinct islands we observe in egg extract (Fig 1A, 2A) conceptually resembles the organization of early Drosophila embryos. During the syncytial blastoderm stage, the cytoplasm of Drosophila embryos is organized into discrete islands of cytoplasm centered on MTOCs, called energids, which function like autonomous cells in many respects (Foe and Alberts, 1983); Foe et al., 2000; Mavrakis et al., 2009). Initially energids travel through less structured bulk cytoplasm to reach the cortex. Later they are kept separate by transient furrows during mitosis, but how they are kept separate during interphase and while migrating, in the absence of plasma membrane barriers, is unclear. Our data shows that AURKB kinase activity localized on microtubule bundles at aster boundaries partitions Xenopus egg cytoplasm into discrete islands. It does so by generating gaps in the actin and keratin networks and by inhibiting microtubule plus end growth that would otherwise blur the boundary. We propose that AURKB, or functionally analogous kinases, may play similar roles in organization of proliferating syncytia.

## Acknowledgements

This work was supported by NIH grant GM39565 (TJM). MBL Fellowships from the Evans Foundation, MBL Associates and the Colwin Fund (T.J.M and C.M.F). Authors thanks the Nikon Imaging Center at Harvard Medical School and Nikon at MBL for imaging support; and the National Xenopus Resource at MBL for Xenopus animals and care.

## Methods and key reagents

### Key Resource Table

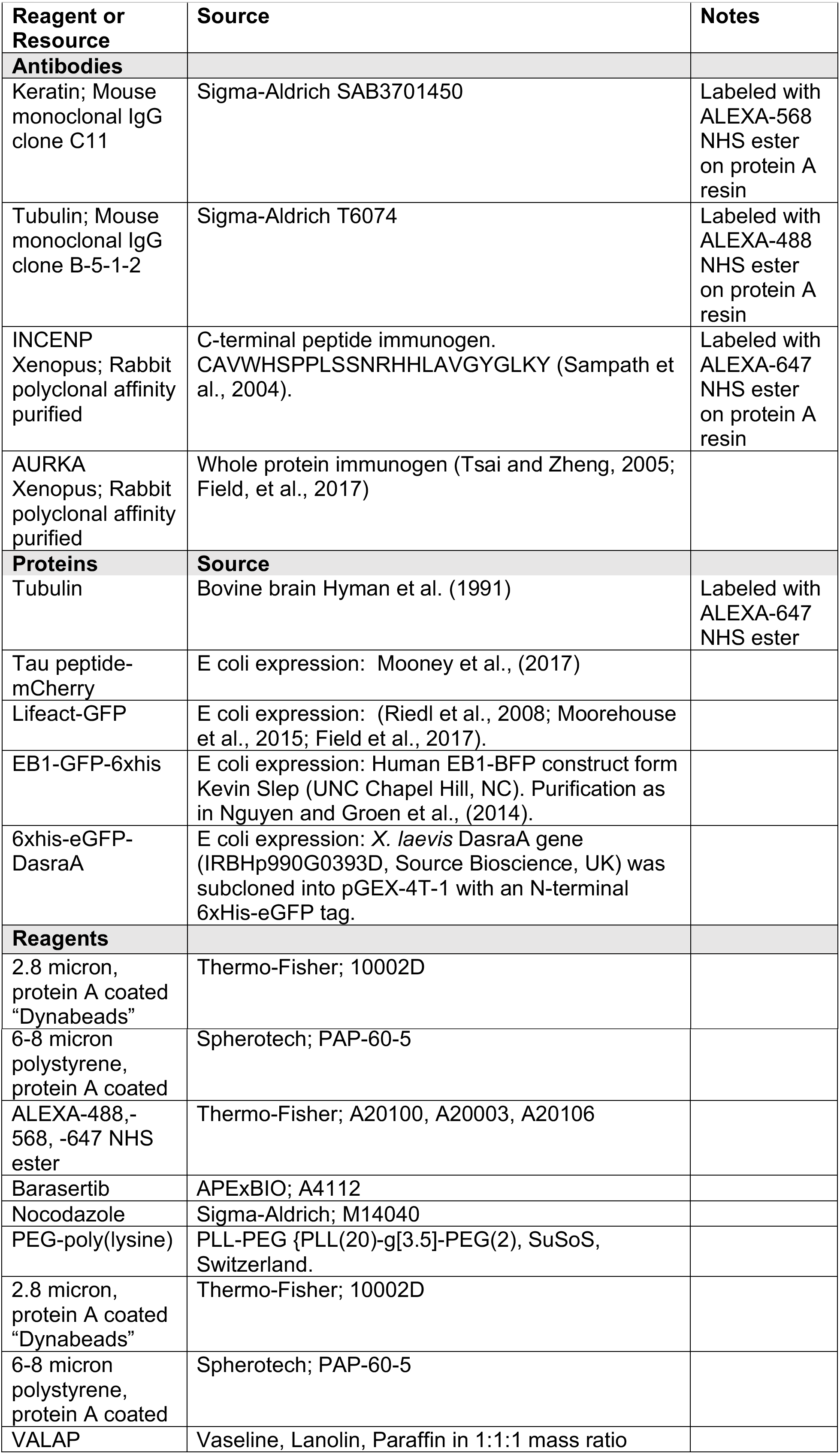

### Xenopus Egg Extracts

Freshly made, actin-intact CSF egg extract was used for all experiments (Field, et al., 2014). To prevent keratin aggregation, we stored extract at 10-12° C, then cooled briefly to 0° C before setting up reactions. Extract stored this way was typically usable for ~ 8 hrs.

#### Interphase aster assembly reactions

Methods were similar to those described previously (Field et al., 2017). In a typical experiment fluorescent probes were added to CSF extract on ice. Calcium chloride was added to 0.4mM final to trigger exit from mitosis, the reaction was mixed well, incubated at 18 °C for 5 min and returned to 0 °C. Anti-AURKA beads were added as MTOCs, and the reaction divided up for drug or vehicle addition. 6.5 μl aliquots were spotted onto the surface of a 22mm^2^ glass coverslip coated with PEG-poly-lysine mounted on a 4-place metal holder. A 18 mm^2^ similarly coated coverslip was immediately place on top. The resulting 20μm thick squash was sealed with molten VALAP and imaging was started immediately. Typically, we imaged 2-4 reactions in parallel using a 20x dry objective at a room temperature of 18-20°C, collecting sequences at 3-4 positions per each reaction for 1 hr. T=0 is the time the reaction was squashed and warmed to RT. Depending on the density of nucleating sites (anti-AURKA beads) asters typically grew into contact at 8-15 min and formed CPC-positive boundaries a few minutes later. Asters and boundaries were stable until ~ 60 min, and eventually decayed with loss of organized microtubules. See Field et., (2017) for more details on assembly reactions.

### Anti-AURKAbeads

(Tsai and Zheng, 2005; Field et al., 2017)

### Passivated Coverslips

Coverslips: 18 × 18 or 22 × 22 mm were coated with poly-lysine PEG (PLL(20)-g{3.5}-PEG(2); SuSOS Chemicals) using a simplified cleaning method. Coverslips were dipped in 70% ethanol, flamed, cooled and placed on a droplet of 100 μg/ml poly-lysine-PEG in 10 mM HEPES, pH 7.4, on parafilm for 15-30 min. They were then washed twice with distilled water for 5 min each, dried with a jet of nitrogen and used the same day.

### Cytoplasmic flow experiments

To induce flow through the squash preparation we waited until CPC-positive boundaries had formed, then placed a 10 gm weight (a bolt) on one side of the coverslip. After 30 sec the bolt was removed and a DIC movie was collected. Fluorescent images were collected before and after inducing flow.

### Anti-INCENP bead experiments

Protein A coated beads (2.8 μm Dynabeads or 6-8 μm polystyrene beads) were saturated with anti-INCENP IgG overnight and then washed with CSF-XB (10 mM K-HEPES, pH 7.7, 100 mM KCl, 1 mM MgCl_2_, 0.1 mM CaCl_2_, 50 mM sucrose and 5 mM EGTA). Beads were added to CSF extract and imaged as per aster assembly reactions above. We typically waited 20-40 min and then imaged multiple fields.

### Quantification of F-actin intensity around anti-INCENP beads

Intensity profiles of Lifeact-GFP signal around anti-INCENP beads were quantified using the Fiji plugin “Radial Profile Extended” by Philippe Carl (https://imagei.nih.gov/ij/plugins/radial-profile-ext.html) with an integration angle of +/- 180 degrees (a full circle). Radial intensity profiles were normalized by dividing by the background intensity, in an unperturbed region far from the bead. The background intensity was estimated as the mean intensity of the last 10 points in each profile, about 80 μm from the bead. As a result, all radial intensity profiles approached the same background value.

### Zygote Immunofluorescence

Embryos were fixed and stained as previously described, Nguyen and Groen et al., (2014). Briefly, embryos were fixed in 50 mM EGTA, ~ pH 6.8, 10% H2O, 90% methanol for 24 hrs at room temperature with gentle shaking. Prior to staining, embryos were rehydrated in a series of steps— 25%, 50%, 75% and 100% TBS (50 mM Tris, pH 7.5, 150 mM NaCl/methanol: 15 min per step with gentle shaking. Embryos were then hemisected in TBS on an agarose cushion using a small piece of razor blade. Embryos were bleached overnight in a solution of 1% H2O2, 5% formamide, 0.5x SSC (75 mM NaCl and 8 mM sodium citrate, pH 7). Embryos were incubated with directly labeled antibodies for at least 24 hours at 4° C with very gentle rotation. Antibodies were diluted in TBSN (10 mM Tris-Cl, pH 7.4, 155 mM NaCl, 1% IGEPAL CA-630), 1% BSA, 2% FCS and 0.1% Sodium Azide. After antibody incubation, embryos were washed in TBSN for at least 48 hr (with several solution changes) then washed 1 X in TBS and 2X in methanol (methanol washes for 20 min each). Embryos were cleared in Murray Clear solution (benzyl benzoate/benzyl alcohol 2:1) and mounted in metal slides (1.2 mm thick). The slides have a hole in the center. The hole was closed by attaching a coverslip to the bottom of the slide using heated parafilm.

## Microscopy

Figures 1 and 2: Reactions were imaging using a 20x Plan Apo 1.4 NA dry objective on a Nikon Ti2 wide field, inverted microscope with a Perfect Focus System using a Nikon DS-Qi2 camera.

Figure 3: Reactions were imaged using a 40x oil immersion objective on a Nikon Ti2 wide field, inverted microscope with a Perfect Focus System using a Hamamatsu Flash4.0 V2+ sCMOS camera.

Figure 4 A,B and G: Reactions were imaged by wide-field fluorescence using a 10x or 20 x Pan Apo 1.4 NA dry objective on a Nikon Ti2 wide field, inverted microscope with a Perfect Focus System using a Hamamatsu Flash4.0 V2+ sCMOS camera.

Figure 4 E, F and H: using a 10x or 20 x Pan Apo 1.4 NA dry objective on an upright Nikon Eclipse 90i microscope with a Hamamatsu ORCA-ER cooled CCD camera.

Figure 5: Fixed Xenopus embryos were imaged using a Nikon Ti-E inverted microscope with a Nikon A1R point scanning confocal head using 10x dry and 20x multi-immersion objectives.

Note that all imaging of extract reactions was performed at 18-20° C.

## Supplementary Movies

Movie 1A F-actin disassembles at boundaries between asters in control egg extract. Disassembly is co-incident with Aurora kinase B localization. F-actin (green) and CPC (red). Corresponds to Figure 1A.

Movie 1B F-actin disassembles at boundaries between asters in control egg extract. Tubulin (cyan) and CPC (red). Corresponds to Figure 1A.

Movie 1C F-actin disassembles at boundaries between asters in control egg extract. Disassembly is co-incident with Aurora kinase B localization. F-actin (green) and CPC (red). Zoom-in of Movie 1A and Figure 1A

Movie 2A F-actin does not disassemble at boundaries between asters in the presence of the AURKB inhibitor barasertib. F-actin (green) and CPC (red). Note the absence of localization of Aurora kinase B between asters and the absence of F-actin disassembly. Corresponds to Figure 1B and is a parallel reaction to Figure 1A.

Movie 2B F-actin does not disassemble at boundaries between asters in the presence of the AURKB inhibitor barasertib. Tubulin (cyan) and CPC (red). Note the absence of localization of Aurora kinase B between asters. Corresponds to Figure 1B and is a parallel reaction to Figure 1A

Movie 3A Keratin disassembles at boundaries between asters in control egg extract. Disassembly is co-incident with Aurora kinase B localization. Keratin (green) and CPC (red). Similar data to the experiment in Figure 2A, different reaction.

Movie 3B Keratin disassembles at boundaries between asters in control egg extract. Disassembly is co-incident with Aurora kinase B localization. Growing MT + ends were visualized via EB1-GFP (cyan) and CPC (red). Similar data to the experiment in Figure 2A, different reaction.

Movie 3C Keratin disassembles at boundaries between asters in control egg extract. Disassembly is co-incident with Aurora kinase B localization. Keratin (green) and CPC (red). In this example, Aurora B kinase localizes late in the reaction. Corresponds to Figure 2B.

Movie 4 Keratin does not disassemble at boundaries between asters in the presence of the AURKB inhibitor barasertib. Keratin (green) and CPC (red). Note the absence of localization of Aurora kinase B between asters and the absence of keratin disassembly. Corresponds to Figure 2C and is a parallel reaction to those in Figure 1A.

Movie 5 Low resistance to pressure induced flow between asters visualized by DIC microscopy. Corresponds to Figure 3C.

Movie 6 Actin, keratin and MT disassembly around CPC beads. Corresponds to Figure 4D.

